# SARS-CoV-2 Protein ORF3a Drives Lysosomal Membrane Atg8ylation

**DOI:** 10.1101/2024.09.12.612614

**Authors:** Deepa Ajnar, Ananya Sarkar, Seema Riyaz, Avanish pandey, Bishnu Prasad Sahoo, Shivam Patnaik, Smruti Ranjan Rout, Poornima Menon, Dipranil Dutta, Pulkit Asati, Priyanka Sharma, Sai Prasad Pydi, Suresh Kumar

## Abstract

Atg8ylation is an autophagy associated response to membrane damage that recruits mammalian ATG8 proteins (mATG8s) to damaged membranes to promote their repair or removal. Here, we show that the SARS-CoV-2 protein ORF3a induces lysosomal membrane atg8ylation and that this response protects cells from death. mATG8s interact with ORF3a and are required for lysosomal damage induced by ORF3a. ORF3a targets mTOR in an Atg8ylation dependent manner and promotes lysophagy through mATG8 mediated coordination of TRIM16 and Galectin-3. ORF3a triggers apoptosis, necroptosis, and pyroptosis, whereas atg8ylation limits ORF3a- induced cell death. Together, our findings identify mATG8s and the autophagy conjugation machinery as key regulators of lysosomal atg8ylation and lysophagy in response to the SARS-CoV-2 virulence factor ORF3a.

## Introduction

Autophagy controls cellular homeostasis by capturing cellular waste and delivering it to lysosomes for degradation. It impacts a wide range of human diseases, making it a potential target for therapies against various human pathologies, including cancer, neurodegenerative diseases, metabolic disorders, infection and inflammation-related diseases, among others(Dikic & Elazar, 2018; Levine & Kroemer, 2019). Autophagy is broadly categorized into canonical and non-canonical. A key distinction is indispensability of ULK1/FIP200/ATG13/ATG101 complex for canonical autophagy, whereas it is not necessary for non-canonical autophagy(Codogno *et al*, 2012). The common link between canonical and non-canonical autophagy is necessity of autophagy conjugation machinery, which conjugates ATG8 proteins to autophagosomal membranes(Mizushima *et al*, 2011).

Atg8ylation involves recruitment of ATG8 proteins to the damaged membranes for their repairs or selective removal (Deretic & Klionsky, 2024; Deretic & Lazarou, 2022; Kumar *et al*, 2021b). In response to membrane stress, atg8ylation recruits endosomal sorting complex required for transport (ESCRT) for repairing the damaged membranes (Javed *et al*, 2023; Jia *et al*, 2022) or autophagy machinery to selectively and efficiently remove damaged membranes (Maejima *et al*, 2013). Atg8ylation is akin to ubiquitination, while ubiquitination tags proteins for degradation, Atg8ylation recognizes damaged membranes and tag them for repair or degradation(Kumar *et al*., 2021b). Similar to ubiquitylation, atg8ylation involves an E1-like activating protein, ATG7 which is conjugated with the C-terminal glycine of Atg8s. This process is followed by the E2-like activity of ATG3 on Atg8 proteins, and subsequently, the conjugation of Atg8 proteins to phosphatidylethanolamine (PE))(Mizushima *et al*., 2011) or phosphatidylserine (PS) (Durgan *et al*, 2021) is regulated by the E3-like complex ATG5-ATG12-ATG16L1 (Kumar *et al*., 2021b). Another similarity between ubiquitin proteasomal system and Atg8ylation is requirement of ubiquitin interacting motifs (UIM)(Hofmann & Falquet, 2001) and ATG8 interacting motifs (AIM)(Morishita & Mizushima, 2019) in the receptors for substrate recognition. Proteins with UIM interact with ATG8s at a Site distinct from the LC3 interacting regions (LIR) docking sites (Marshall *et al*, 2019). Therefore, both ubiquitination and Atg8ylation are in principle similar processes that handle different cargo(Kumar *et al*., 2021b). Conjugation of ATG8s to single membrane (CASM)(Goodwin *et al*, 2021) is a process similar to atg8ylation, however, atg8ylation can happen during canonical autophagy where ATG8s can be recruited to double membraned autophagosomes during stress(Kumar *et al*., 2021b), whereas, CASM exclusively involves conjugation of ATG8 proteins to single membranes(Durgan & Florey, 2022).

Lysosomal damage induces Atg8ylation which recruits ESCRTs machinery for repair or autophagy machinery for degradation and removal of damaged lysosomes by lysophagy(Skowyra *et al*, 2018). Galectin 3 (Gal3) was believed to sense the lysosomal damage and tag the damaged lysosomes for autophagy mediated degradation(Maejima *et al*., 2013; Skowyra *et al*., 2018). However, more recent studies have shown that Gal3 not only tag the damaged lysosomes for autophagy mediated degradation, it also helps recruiting ESCRTs machinery for repairing the membranes (Jia *et al*, 2020).

Canonical autophagy is controlled by mammalian target of rapamycin (mTOR) and AMPK, where mTOR inhibits autophagy and AMPK activates autophagy by phosphorylating ULK1 at different sites(Kim *et al*, 2011). mTOR also inhibits autophagy by phosphorylating transcription factor EB and preventing its nuclear translocation(Settembre *et al*, 2011; Settembre *et al*, 2012). mATG8s and autophagy conjugation machinery have been previously reported to facilitate nuclear translocation of TFEB by inhibiting mTOR(Kumar *et al*, 2020a; Nakamura *et al*, 2020).

SARS-CoV-2 infection and its non-structural proteins and accessory factors target autophagy(Chen *et al*, 2021; Kumar *et al*, 2021a). ORF3a has been shown to inflict lysosomal damage (Miao *et al*, 2021; Walia *et al*, 2024). SARS-CoV-2 ORF3a interacts with components of endolysosomal system, and potentially inhibit their cellular function to facilitate viral egress(Chen *et al*., 2021; Ghosh *et al*, 2020; Miao *et al*., 2021).

Our study demonstrates that expression of SARS-CoV-2 factor ORF3a induces Atg8ylation of lysosomal membranes. mATG8s interact with ORF3a and are required for lysophagy induced by ORF3a. ORF3a induces lysophagy by organizing TRIM16 complexes with galectin3 and mATG8s are required for ORF3a induced lysophagy. mATG8s are required for ORF3a mediated mTOR inhibition consequent TFEB activation. Finally, mATG8s selectively remove damaged lysosomes and protect cells from panoptosis induced by ORF3a.

## Results

### SARS-CoV-2 factor ORF3a induces Atg8ylation by mTOR inhibition

SARS-CoV-2 protein ORF3a has been reported to induce lysosomal damage(Miao *et al*., 2021; Walia *et al*., 2024) and other lysosomal related functions including lysosomal exocytosis(Chen *et al*., 2021; Ghosh *et al*., 2020; Miao *et al*., 2021). Consistent with previous reports we observed impaired lysosomal quality reflected by reduced lysotracker intensity in ORF3a-mCherrry expressing cells (Figure 1A,B), in contrast, expression of SARS-CoV-2 non-structural proteins NSP3, NSP4 and NSP6 did not alter lysosomal quality (Figure S1A,B). Recent studies have shown the role of mammalian ATG8 proteins (mATG8) proteins in lysosomal function(Goodwin *et al*., 2021; Gu *et al*, 2019; Kumar *et al*., 2020a; Kumar *et al*, 2020b; Nakamura *et al*., 2020). We wondered whether mATG8s participate in lysosomal damage induced by ORF3a. To test this, we used previously characterized Hexa^KO^ cells, where six mATG8s (LC3A, LC3B, LC3C, GABARAP, GABARAPL1 and GABARAPL2) are knocked out (Nguyen *et al*, 2016). We observed reduced effect of ORF3a mediated lysosomal stress in Hexa^KO^ cells (Figure 1A,B). ORF3a-mCherry co-localized with lysosomal damage sensor galectin 3 (Gal3)(Maejima *et al*., 2013), colocalization between Gal3 and ORF3a was reduced in Hexa^KO^ cells (Figures 1 C,D and S1C,D). Endogenous and GFP-Gal3 puncta formation was increased after ORF3a expression in HeLa^WT^ (wild type HeLa cells) but not in Hexa^KO^ cells (Figures 1C,D and S1E,F). ORF3a expression resulted in increased localization of Gal3 to lysosomes in HeLa^WT^ but not in Hexa^KO^ cells (Figure 1E,F). Lysosomal stress influences nuclear translocation of TFEB (transcription factor EB)(Chauhan *et al*, 2016; Jia *et al*, 2018; Nakamura *et al*., 2020), ORF3a expression resulted nuclear translocation of endogenous and exogenously expressed GFP-TFEB (Figure 1G, H and S1 G,H) which was reduced in Hexa^KO^ cells (Figures 1G, H). Lysosomal damage inhibits mammalian target of rapamycin (mTOR) function(Chauhan *et al*., 2016). We employed western blotting to quantify the effect of ORF3a expression on well-known mTOR substrate P70S6K (Sancak *et al*, 2008). ORF3a expression inhibited P70S6K phosphorylation in HeLa^WT^ cells but not in Hexa^KO^ cells (Figures 1I, J). Lysosomal damage causes displacement of mTOR from lysosomes (Zoncu *et al*, 2011). ORF3a expression reduced mTOR localization to lysosome in HeLa^WT^ cells, however, this effect was significantly reduced in Hexa^KO^ cells (Figure 1 K, L and S1I). TFEB is a substrate of mTOR, TFEB and mTOR interact on lysosomes (Settembre *et al*., 2012), since ORF3a expression affected nuclear translocation of TFEB and inhibits mTOR, we wondered whether ORF3a affects TFEB and mTOR complexes. Interaction of TFEB with mTOR was abolished in in ORF3a expressing HeLa^WT^ cells but not in Hexa^KO^ cells (Figure 1M,N). To test whether ORF3a affects lysosomal function by inhibiting mTOR, we used mutant of RagB (RagB^Q99L^), which keeps mTOR in constitutively active state (Zoncu *et al*., 2011). Expression of RagB^Q99L^ prevented ORF3a induced lysosomal damage as reflected by lysotracker (Figure 1 O and S2A) and Gal3 puncta formation (Figure 1P and S2B). Next, we depleted TSC2 (Tuberous sclerosis 2) using CRISPR CAS9 system. TSC2 inhibits mTOR(Inoki *et al*, 2002; Inoki *et al*, 2003), and depletion of TSC2 activates mTOR. We expressed ORF3a in HeLa^WT^ or TSC2 depleted cells, ORF3a expression resulted in lysosomal damage as shown by increase in Gal3 puncta in HeLa^WT^ but not in TSC2 depleted cells (Figures 1Q and S2 C). Thus, ORF3a induces Atg8ylation of lysosomal membranes by inhibiting mTOR.

**Figure 1.**
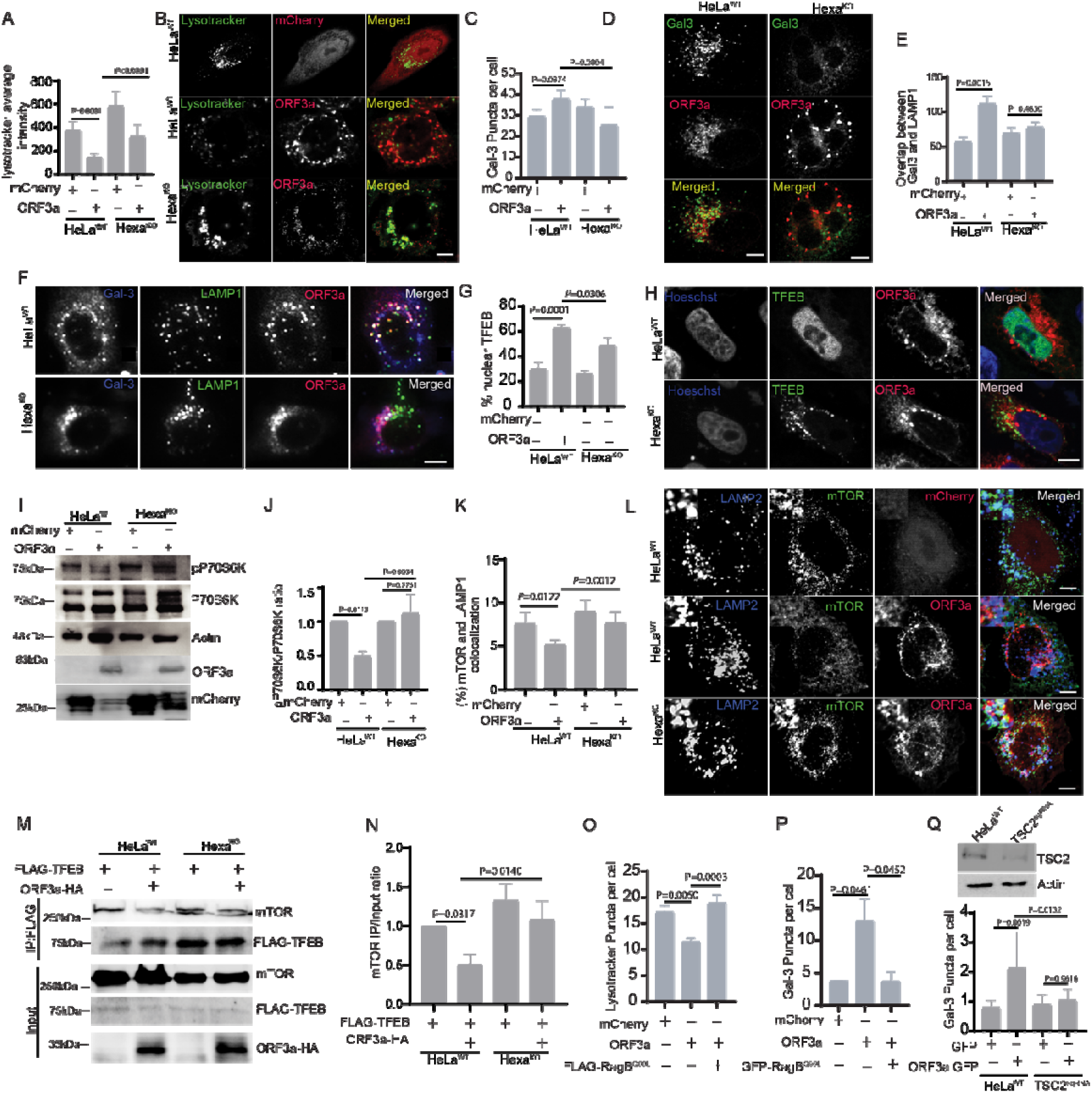
ORF3a inflicts Atg8ylation of lysosomal membrane. **(A,B)** High content microscopy (HCM) quantification and confocal microscopy analysis of the effect of Hexa^KO^ on lysotracker intensity in ORF3a expressing cells (>500 primary objects/cells examined per well; minimum number of wells, 5). p values were determined using GraphPad prism (n=9) ANOVA. Scale bar 5 µm. **(C,D)** HCM quantification and confocal microscopy analysis of the effect of Hexa^KO^ on ORF3a induced Gal3 puncta formation. p values were determined using GraphPad prism (n=8) ANOVA. Scale bar 5 µm. **(E)** HCM analysis of the effect of ORF3a on Gal3 and LAMP1 colocalization in HeLa^WT^ and Hexa^KO^ cells. (>500 primary objects examined per well; minimum number of wells, 6). p values were determined using GraphPad prism (n=6) ANOVA. **(F)** Representative micrographs showing the effect of ORF3a expression on Gal3 and LAMP1 colocalization in HeLa^WT^ and Hexa^KO^ cells. Scale bar 5 µm. Pseudo colours: Red, ORF3a; Blue, Gal3 and Green, LAMP1. **(G,H)** HCM and confocal image analysis of effect of Hexa^KO^ on ORF3a-mCherry induced nuclear translocation of TFEB (>500 primary objects/cells examined per well; minimum number of wells,10). p values were determined using GraphPad prism (n=10) ANOVA. Scale bar 10 µm**. (I, J)** Western blot analysis of the effect of ORF3a- mCherry expression on mTOR substrate p70S6K and TFEB, in HeLa^WT^ and Hexa^KO^ cells and a graph showing quantification of pP70S6K normalized with total P70S6K in Hela^WT^ and Hexa^KO^ cells. **(K,L)** HCM quantifications and confocal image analysis of the effect of ORF3a-mCherry expression on colocalization between mTOR and LAMP1 in HeLa^WT^ and Hexa^KO^ cells. p values were determined using GraphPad prism (n=6) ANOVA; HCM, >500 cells counted per well; minimum number of valid wells 6. Scale bar 5 µm **(M)** Co-IP analysis of the effect of ORF3a expression on interaction between FLAG-TFEB and endogenous mTOR in HeLa^WT^ and Hexa^KO^ cells. **(N)** A graph shows quantification of effect of ORF3a-HA expression on interaction between TFEB an mTOR in WT and Hexa^KO^ cells normalized with inputs (n=3). **(O)** HCM quantification of the effect of RagB^Q99L^ expression on ORF3a mediated inhibition of lysotracker puncta. Data, means ± SEM, p values were determined using GraphPad prism (n=6) ANOVA; HCM, >500 cells counted per well; minimum number of valid wells 6. **(P)** HCM quantification of the effect of RagB^Q99L^ expression on ORF3a mediated formation of Gal3 puncta. Data, means ± SEM, p values were determined using GraphPad prism (n=6) ANOVA; HCM, >500 cells counted per well; minimum number of valid wells 6. **(Q)** Immunoblot showing knock out of TSC2 in HeLa cells. HCM quantification of effect of ORF3a expression on Gal3 puncta formation in HeLa^WT^ and TSC2^KO^ cells. p values were determined using GraphPad prism (n=5) ANOVA.

### ORF3a interacts with mATG8s

Our data indicated the role of mATG8 proteins in ORF3a induced lysosomal damage. We asked if ORF3a expression influences recruitment of ATG8s to lysosomes. ORF3a expression resulted in increased puncta formation of all mATG8s except GFP-LC3A (Figure 2 A,B and S2D). In a co-immunoprecipitation (co-IP) experiment, ORF3a-HA interacted with GFP tagged LC3B, LC3C, GABARAP and GABARAPL1 (Figures 2C). Consistent with co-IP data, ORF3a-mCherry colocalized with all mATG8s. Strongest colocalization was seen with GFP-LC3B followed by GFP-GABARAP, GFP-GABARAPL2, GFP-LC3C, GFP-GABARPL1 and GFP-LC3A (Figure S2 E,F). Next, we tested whether ORF3a influences mATG8s puncta formation at endogenous levels. We observed increased puncta formation in LC3B (Figure S2 G,H), LC3C and GABARAP (Figure 2D,E). Similar to GFP tagged mATG8s, endogenous LC3B, LC3C and GABARAP colocalized with ORF3a- mCherry (Figure 2G and S2I). Moreover, localization of GFP labelled GABARAPs to lysosomes was increased in ORF3a expressing cells (Figure S2 J,K). Keeping up with these data, ORF3a recruited endogenous LC3B, LC3C and GABARAP to lysosomes (Figure 2F,G). However, GABARAP recruitment to lysosome was > 3-fold higher than that of LC3B and LC3C (Figure 2F,G). Consistent with this data, GFP- GABARAP and GFP-GABARAPL1 rescued ORF3a induced Gal3 puncta formation in HexaKO (Figure 2 H,I). Keeping up with these data, we observed an increase in Gal3 localization to lysosomes in Hexa^Ko^ cells rescued with GFP-GABARAP (Figure S2 L,M). ORF3a contains a YXXΦ based sorting motif (160–163 amino acids)(Ren *et al*, 2020), mutation in this motif reduces the effect of ORF3a on lysosomal damage and autophagy induction (Walia *et al*., 2024). We tested whether expression of ORF3a^160Y-A/163L-G^ mutant has any effect on Gal3 recruitment to lysosomes. Gal3 puncta formation was reduced in ORF3a^160Y-A/163L-G^ mutant (Figure S2 N,O), consistent with this, localization of Gal3 to lysosomes was reduced in ORF3a-160Y- A/163L-G mutant (Figure S2 P,Q). These data indicate that ORF3a requires autophagy conjugation machinery for inducing lysosomal damage.

**Figure 2.**
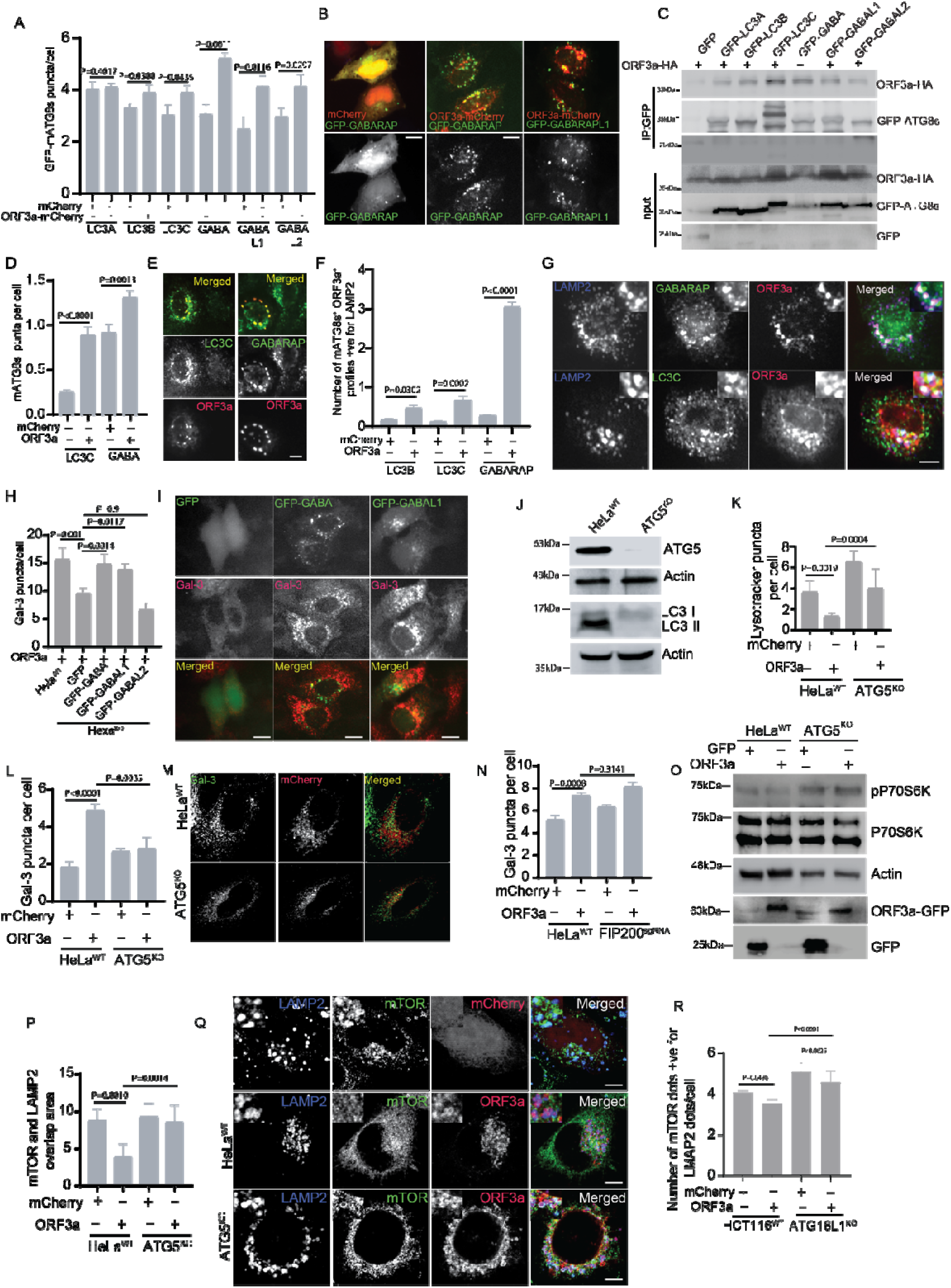
ORF3a induces Atg8ylation by interacting with mATG8s and it is independent of autophagy initiation machinery. **(A)** HCM quantification of the effect of ORF3a expression on mATG8s dot formation. GFP tagged LC3A, LC3B, LC3C, GABARAP, GABARAPL1 and GABARAPL2 plasmids were co-transfected with mCherry or ORF3a-mCherry into HeLa cells. GFP-mATG8 puncta were counted using computer defined algorithm. (>500 primary objects examined per well; minimum number of wells, 5). p values were determined using GraphPad prism (n=5) ANOVA. **(B)** Fluorescence microscopy analysis of the effect of mCherry or ORF3a-mCherry expression on GFP-GABARAP and GFP-GABARAPL1 dot formation. Scale bar 5 µm. **(C)** Co-IP analysis of interactions between ORF3a-HA and GFP-LC3A, GFP-LC3B, GFP-LC3C, GFP-GABARAP, GFP-GABARAPL1 and GFP-GABARAPL2 in 293T cells. **(D,E)** HCM quantification and micrograph analysis of colocalization between ORF3a with LC3C and GABARAP in HeLa cells. GFP tagged LC3C and GABARAP plasmids were co-transfected with ORF3a-mCherry into HeLa cells, GFP and mCherry transfected cells were gated and used for quantification (>500 primary objects examined per well; minimum number of wells, 8). p values were determined using GraphPad prism (n=8) ANOVA. Scale bar 5 µm. **(F,G)** HCM quantification and image analysis of triple colocalization between ORF3a- mCherry with LC3B, LC3C, GABARAP and LAMP2. p values were determined using GraphPad prism (n=5) ANOVA; scale bar 5µm. **(H,I)** HCM quantification and micrograph analysis of the effect of complementation of Hexa^KO^ HeLa cells with GFP, GFP-GABARAP, GFP-GABARAPL1 and GFP-GABARAPL2 on formation of endogenous Gal3 dots in response to ORF3a-HA expression. (>500 primary objects examined per well; minimum number of wells, 6). Scale bar 5 µm. **(J)** An immunoblot showing knock out of ATG5 in HeLa cells. **(K)**HCM quantification analysis of the effect of ATG5^KO^ on lysotracker intensity in ORF3a-mCherry expressing HeLa cells. **(L,M)** HCM quantification and confocal image analysis of the effect of ATG5^KO^ on Gal3 puncta formation in HeLa cells expressing mCherry or ORF3a-mCherry. **(N)** HCM quantification of the effect of FIP200^KO^ on Gal3 puncta formation in ORF3a- mCherry expressing HeLa cells. **(O)** An immunoblot analysis of effect of ORF3a expression on mTOR substrate pP70S6K in HeLa^WT^ and ATG5^KO^ cells. **(P)** HCM quantification analysis of the effect of ATG5^KO^ on colocalization between mTOR and LAMP2 to assess localization of mTOR to lysosomes in ORF3a-mCherry expressing HeLa cells. **(Q)** Micrograph showing the effect of ORF3a-mCherry expression on mTOR and LAMP2 colocalization in HeLa^WT^ or ATG5^KO^ cells. Scale bar 5 µm. **(R)** HCM quantification to analyze the effect ORF3a-mCherry expression on colocalization between mTOR and LAMP2 in HCT116^WT^ and ATG16L^KO^ cells.

### ORF3a induced Atg8ylation is dependent on ATG8 conjugation machinery

Atg8ylation involves recruitment of mATG8 proteins to damaged membranes and is dependent on ATG conjugation machinery. To examine whether conjugation machinery is involved in ORF3a induced Atg8ylation of lysosomal membranes, we generated ATG5^KO^ and ATG16L1^KO^ using CRISPR/CAS9 system (Figures 2J and S3A). We employed HCM to quantify the effect of ORF3a expression on lysotracker intensity. We observed reduced lysotracker intensity in HeLa^WT^ cell but not in ATG5^KO^ and ATG16L1^KO^ cells expressing ORF3a (Figures 2K S3C). Conversely, CRISPR mediated depletion FIP200 (Figure S3D), a component of autophagy initiation complex(Hara *et al*, 2008; Hosokawa *et al*, 2009), did not impact ORF3a mediated change in lysotracker intensity (Figure S3D,E). Furthermore, ORF3a induced Gal3 puncta formation was reduced in ATG5^KO^ cells (Figure 2L,M) but not in FIP200 downregulated cells (Figures 2N and S3G). Consistently, ORF3a expression enhanced Gal3 localization to lysosomes in HeLa^WT^ cells but not in ATG5^KO^ cells (Figure S3H,I). However, FIP200 depletion did not affect ORF3a induced localization of Gal3 to lysosomes (Figure S3J,K). Next, we tested whether autophagy conjugation machinery is required for ORF3a mediated mTOR inhibition. ORF3a expression resulted in reduced P70S6K phosphorylation in HeLa WT cells; however, no such effect was observed in ATG5^KO^ cells (Figure 2O). Furthermore, ORF3a inhibited mTOR localization to lysosome was reduced in wild type cells but not in ATG5^KO^ and ATG16L1KO cells (Figures 2 K-M and S3L). In contrast, FIP200 depletion could not avert the effect of ORF3a expression on P70S6K phosphorylation (Figures S3M). In addition, FIP200 downregulation did not alter the effect of ORF3a on displacing mTOR from lysosomes (Figure S3 N, O). Thus, ORF3a induces Atg8ylation of lysosomal membranes.

### mATG8 facilitate the recruitment of TRIM16 and Galectin-3 complexes to damaged lysosomes

Our data indicate that ORF3a induces lysosomal damage and mATG8s facilitate autophagy mediated selective removal of damaged lysosomes. Previous studies have shown that ORF3a interacts with lysophagy receptor TRIM16 and lysophagy sensor Gal3(Stukalov, 2020). TRIM16 interacts with mATG8s (Chauhan *et al*., 2016; Mandell *et al*, 2014). Based on this, we asked if mATG8s have role in TRIM16 and ORF3a associations. Using HCM we examined the colocalization of TRIM16 with ORF3a, we found colocalization of FLAG-TRIM16 with ORF3a-mCherry, this colocalization was reduced in HexaKO cells (Figure 3A and S3P). ORF3a mediated TRIM16 interacts with Gal3 to orchestrate lysophagy(Chauhan *et al*., 2016). We observed reduced colocalization between Gal3 and TRIM16 in ORF3a expressing HexaKO cells compared to wild type cells (Figure 3 B,C). Consistent with these data, Interactions of TRIM16 with ORF3a and Gal3 were reduced in HexaKO cells (Figure 3D). These data indicate that mATG8s organize TRIM16-Gal3 complexes in response to ORF3a induced lysosomal damage.

**Figure 3.**
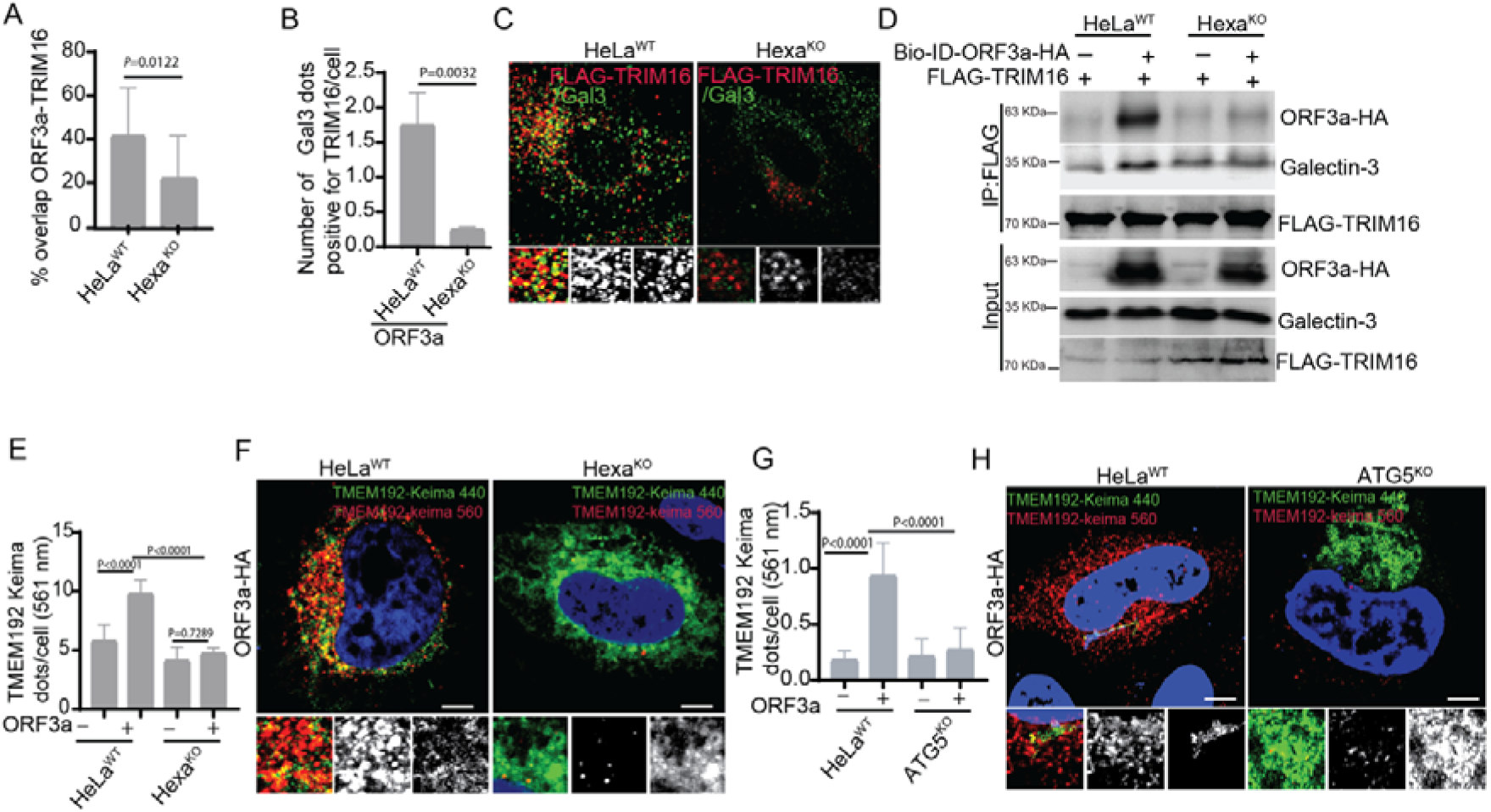
mATG8s regulates ORF3a induced lysophagy. **(A)** HCM quantification analysis of the effect of Hexa^KO^ on colocalization between ORF3a-mCherry and FLAG-TRIM16 in HeLa cells. p values were determined using GraphPad prism (n=6) t test. **(B,C)** HCM quantification and confocal image analysis of the effect of Hexa^KO^ on ORF3a induced colocalization between FLAG-TRIM16 and Gal-3 in HeLa cells. p values were determined using GraphPad prism (n=6) t test. **(D)** Co-IP analysis of the effect of Hexa^KO^ on interactions of FLAG-TRIM16 with ORF3a-HA and Galectin-3 in HeLa cells. **(E,F)** HCM quantification and image analysis of effect of Hexa^KO^ on ORF3a induced lysophagy. HeLa^WT^ and Hexa^KO^ cells were transfected with ORF3a- HA and TMEM192-Keima, fluorescence at 561nm and 440nm was quantified in HCM. Scale bar 5µm. **(G,H)** HCM quantification and image analysis of effect of ATG5^KO^ on ORF3a induced lysophagy. WT and ATG5^KO^ cells were transfected with ORF3a-HA aand TMEM192-Keima, fluorescence at 561nm and 440nm was quantified in HCM. Scale bar 5µm.

### mATG8s and ATG5 are required for ORF3a induced lysophagy flux

We next sought to measure lysophagy flux induced by ORF3a using Keima probes in wild type cells and compare with Hexa^KO^ cells. To test this, we employed a recently reported TMEM-192-Keima probe, which specifically measures lysophagy flux (Shima *et al*, 2023). Keima goes through a spectral shift from 440nm to 620nm under lysosomes pH, making it a convenient probe for measuring lysosomal mediated degradation(Katayama *et al*, 2011). Our HCM and confocal microscopy data suggested ORF3a mediated lysophagy flux in HeLa^WT^ cells, however, ORF3a expressing HexaKO or ATg5KO cells did not show lysophagy flux in (Figure 3 E-H). Thus, mATG8s are required for lysophagy induced by ORF3a.

### mATG8s control the expression of genes involved in lysosomal function

mATG8s transcriptionally regulate the function of genes involved in lysosomal function(Kumar *et al*., 2020a). We performed RNA-seq in HeLa^WT^ and Hexa^KO^ cells expressing ORF3a. RNA-seq detected upregulation of 576 genes and downregulation of 530 genes in Hexa^KO^ cells compared to HeLa^WT^ cells expressing ORF3a (Figure S4A, S4B, S4C Table S1). Expression of lysosomal hydrolases e.g., CTSZ, CTSB, HYAL1, PSAP was reduced in Hexa^KO^ (Figure 4A and Table S1). Surprisingly, expression of ATP6V0E1, ATP6V1C1 and ATP6V0A2 that maintain lysosomal acidification was increased (Figure 4A). This partially explained the increased lysotracker intensity in ORF3a expressing Hexa^KO^ (Figure 1 A,B). A recent study reported the role of autophagy receptors NDP52 and OPTN in lysophagy(Shima *et al*., 2023). RNA-seq detected downregulation of NDP52 (CALCOCO2) and optineurin (OPTN) in Hexa^KO^ cells expressing ORF3a (Figure 4A and Table S1). In addition, the expression of autophagy related genes ATG2A, EPG5, VMP1, FYCO1 and ULK1 was downregulated in Hexa^KO^ cells (Figure 4A). Expression of negative regulators of mTOR, DEPTOR(Peterson *et al*, 2009) and DDIT4(Brugarolas *et al*, 2004) was reduced in Hexa^KO^ cells. These data were consistent with the data shown in Figures 1 J-M.

**Figure 4.**
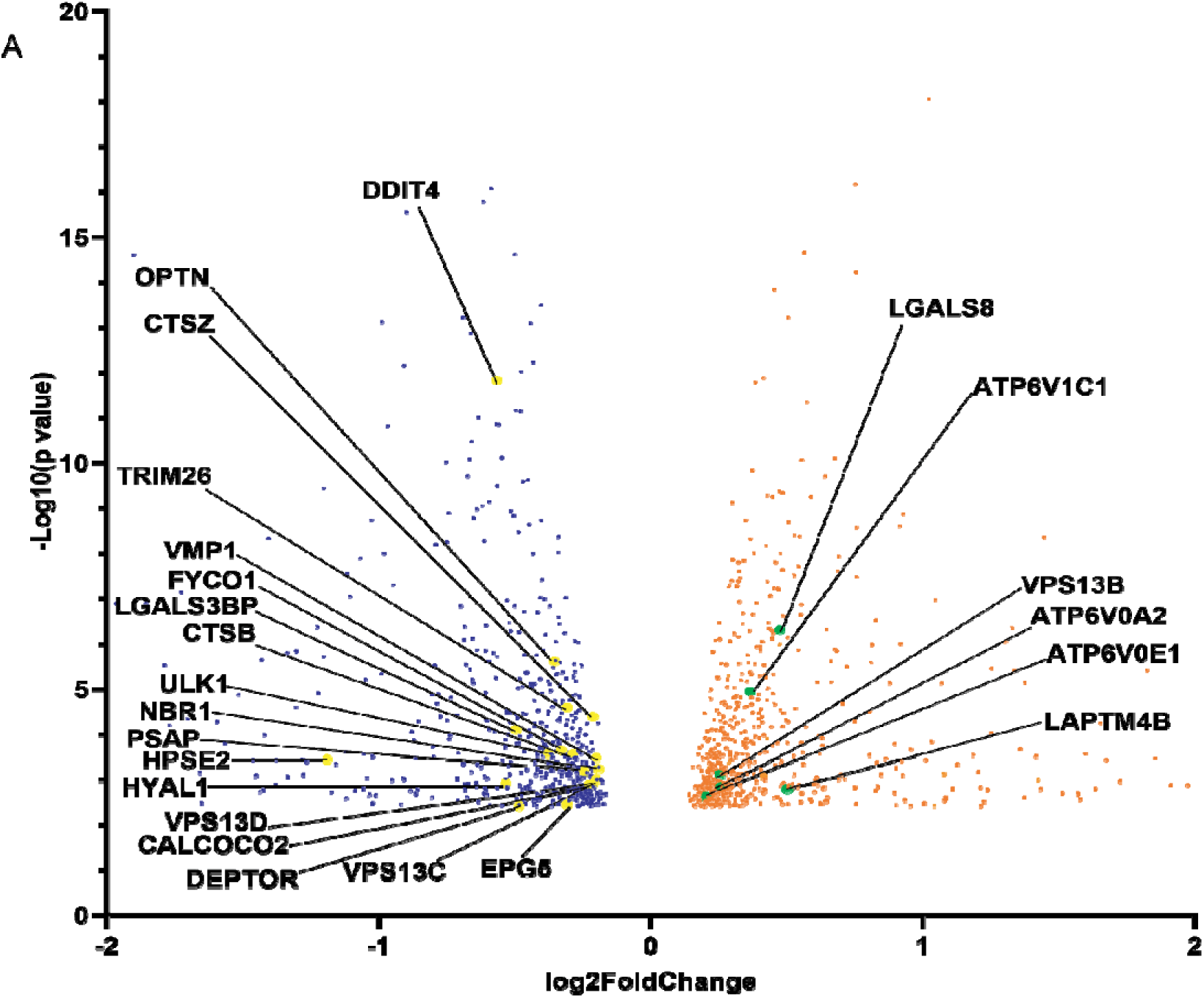
mATG8s affect expression of genes involved lysosomal function. **(A)** Volcano plot showing the effect of Hexa^KO^ on differential gene expression (log2 (fold change); ratio Hexa^KO^/HeLa^WT^). Named genes are the lysosomal hydrolases or genes involved in autophagy and lysosomal function. Yellow circles indicate genes downregulated in Hexa^KO^ cells; green circles genes upregulated in Hexa^KO^ cells. Cut-off used in the plot (P < 0.05). *P* values were calculated using Fisher’s exact test adapted for over dispersed data; edgeR models read counts with NB distribution (see Methods for more). n = 3 biologically independent experiments.

### mATG8s and ATG conjugation machinery protect against the cell death induced by ORF3a expression

SARS-CoV-2 infection or expression of its protein ORF3a induces cell death (Zhang *et al*, 2024; Zhang *et al*, 2022). We asked if ATg8ylation protects against the cell death induced by ORF3a expression. To examine this, we employed HCM to measure propidium iodide (PI) uptake (Claude-Taupin *et al*, 2021). ORF3ainduced PI uptake was elevated in Hexa^KO^ cells (Figure 5A,B). mATG8s comprises of LC3 subfamily (LC3A, LC3B, LC3C) and GABARAP subfamily (GABARAP, GABARAPL1 and GABARAPL2). Out of these two subfamilies, only GABARAP subfamily has been reported to controls major process related to autophagy including autophagosome and lysosome fusion(Nguyen *et al*., 2016), nuclear translocation of TFEB(Goodwin *et al*., 2021; Kumar *et al*., 2020a; Nakamura *et al*., 2020) and lysosomal quality control(Gu *et al*., 2019). Surprisingly, we found that both GABARAP and LC3 subfamilies of mATG8s were required for cell death induced by ORF3a, as ORF3a induced PI uptake was increased in LC3^TKO^ (where LC3A, LC3B and LC3C were knocked out; Figure 5C,D) as well as in GABARAP^TKO^(where GABARAP, GABARAPL1 and GABARAPL2 were knocked out; Figure 5 E,F)(Nguyen *et al*., 2016). We next tested the role of conjugation machinery in ORF3a induced cell death. PI uptake in response to ORF3a expression was increased in ATG16L1^KO^ and ATG5^KO^ cells (Figure S5 A-C).

**Figure 5.**
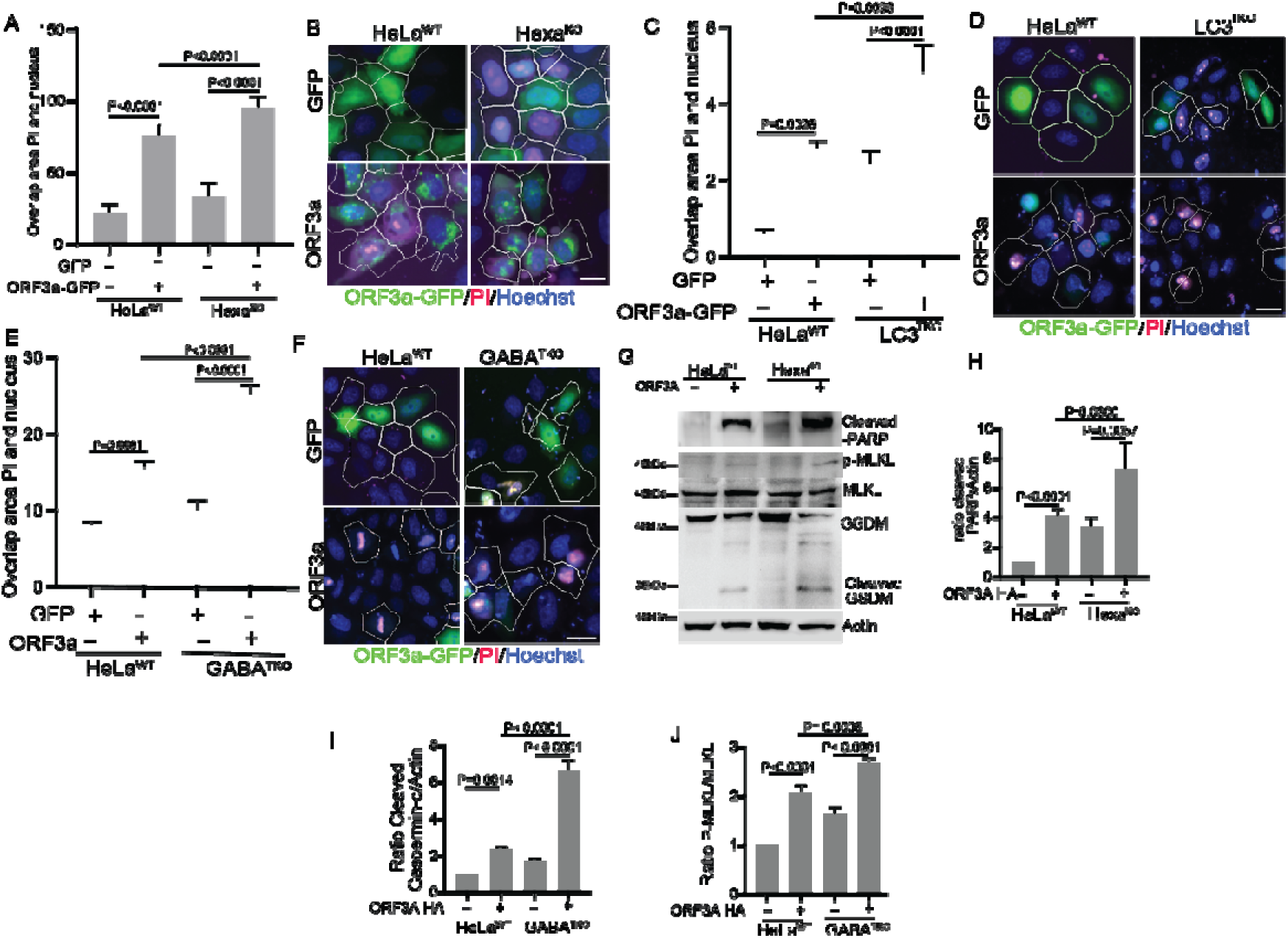
Atg8ylation protects against ORF3a induced cell death. **(A,B)** HCM quantification and image analysis of the effect of Hexa^KO^ on ORF3a mediated colocalization of PI and Hoechst. Overlap between PI and Hoechst was quantified in GFP or ORF3a-GFP transfected cells. Masks; white: GFP or ORF3a-GFP expressing cells, algorithm-defined cell boundaries, Red; Propidium Iodide. Scale bar 10 µm. WT and Hexa^KO^ HeLa cells were transfected with GFP or ORF3a-GFP, cells were stained with PI and analysed under HCM for PI incorporation assay. Masks; white: GFP or ORF3a-GFP expressing cells, algorithm-defined cell boundaries. p values were determine using GraphPad prism (n=9) ANOVA. Scale bar 5 µm. **(C,D)** WT and LC3^TKO^ HeLa cells were transfected with GFP or ORF3a- GFP, cells were stained with PI and analysed under HCM for PI incorporation assay. Colocalization of PI and Hoechst stain was quantified in GFP or ORF3a-GFP transfected cells. Masks; white: GFP or ORF3a-GFP expressing cells, algorithm- defined cell boundaries. p values were determine using GraphPad prism (n=9) ANOVA. Scale bar 10 µm. **(E,F)** WT and GABA^TKO^ HeLa cells were transfected with GFP or ORF3a-GFP, cells were stained with PI and analyzed under HCM for PI incorporation assay. Colocalization of PI and Hoechst stain was quantified in GFP or ORF3a-GFP transfected cells. Masks; white: GFP or ORF3a-GFP expressing cells, algorithm-defined cell boundaries. p values were determine using GraphPad prism (n=9) ANOVA. Scale bar 10 µm. **(G)** Immunoblot analysis of the effect of Hexa^KO^ on ORF3a induced cleavage of PARP-1, Gasdermin-d and on level of p-MLKL in HeLa cells. **(H)** A graph shows the quantification cleaved PARP- normalized with total MLKL in ORF3a expressing WT and Hexa^KO^ HeLa cells (n=3). **(L)** A graph shows the quantification of cleaved Gasdermin-d normalized with actin in ORF3a expressing WT and Hexa^KO^ HeLa cells (n=3). **(I)** A graph shows the quantification of p-MLKL normalized with total MLKL in ORF3a expressing WT and Hexa^KO^ HeLa cells (n=3). **(J)** A model showing the effect of ORF3a expression on lysosomes and consequent Atg8ylation in wild type cells (left) and increased cell death in ATG8s knock-out cells because of the absence of Atg8ylation (right).

The cell viability in ORF3a expressing cells was reduced Hexa^KO^, GABARAP^TKO^ and ATG5^KO^ cells (Figure S5 D-F. Interestingly, FIP200 depletion did not affect ORF3a induced PI uptake (Figure S5 G,H) and cell viability (Figure S5 I). ORF3a contains a YXXΦ based sorting motif (160–163 amino acids), mutation in this motif reduces the effect of ORF3a on lysosomal quality and its effect on autophagy flux(Ren *et al*., 2020; Walia *et al*., 2024). PI uptake induced by ORF3a was significantly reduced in cells expressing Y160A/V163G mutant of ORF3a (Figure S5 J,K). Thus, ORF3a expression inflicts apoptosis due to lysosomal damage (Boya *et al*, 2003; Wang *et al*, 2018) and ORF3a induced cell death is counteracted by Atg8ylation and FIP200 is not required for Atg8ylation.

### Atg8ylation plays a protective role against panoptosis induced by ORF3a

SARS-CoV-2 infection and ORF3a expression inflict different types of cell deaths (Ferren *et al*, 2021; Ren *et al*., 2020; Yuan *et al*, 2023). Recently a new type of inflammation induced cell death, panoptosis, was discovered which displays the features of apoptosis as well as pyroptosis(Gao *et al*, 2024; Lee *et al*, 2021; Pandian & Kanneganti, 2022). We observed increased PARP-cleavage (apoptosis), gasdermin D cleavage (pyroptosis) and increased phosphorylation of MLKL (necroptosis) in ORF3a expression cells (Figure 5 G). These data indicate that ORF3a induces panoptosis. Furthermore, we observed increased PARP-1 cleavage, gasdermin D cleavage and phosphorylation of MLKL in ORF3a expressing Hexa^KO^ cells (Figure 5 H-J), indicating the role of mATG8s in protecting against ORF3a induced panoptosis. Thus, mATG8s control ORF3a induced Atg8ylation which in turn protects against cell death.

## Discussion

In this study, we identify a role for the SARS-CoV-2 accessory protein ORF3a in membrane Atg8ylation and elucidate the mechanism underlying this response. We demonstrate that ORF3a expression induce Atg8ylation of lysosomal membranes. Although ORF3a has previously been shown to cause lysosomal damage and impair autophagosomal fusion with lysosomes, the mechanisms by which it disrupts the autophagy and lysosomal pathway remain poorly understood. Our findings reveal that ORF3a interacts with mATG8 proteins, which promote lysophagy by coordinating the assembly of TRIM16 and Gal-3 complexes. We further show that ORF3a expression suppress mTOR activity in an mATG8 dependent manner. Notably, ORF3a induced lysosomal damage requires the ATG8 conjugation machinery, whereas the ULK1 complex is dispensable for ORF3a-induced lysophagy. Finally, ORF3a induces panoptosis, characterized by the activation of multiple cell death pathways, and this cytotoxicity is substantially limited by mATG8s and components of the ATG8 conjugation machinery. Together, our findings establish a mechanistic link between ORF3a induced lysosomal damage, Atg8ylation, and cell death, revealing an important role for the mATG8 system in the cellular response to SARS-CoV-2 infection.

Atg8ylation is cellular response to membrane stress, wherein ATG conjugation machinery and potentially other related factors such as TECPR1 recruit ATG8 proteins to damage membranes which in turn recruit cellular repair machinery, ESCRTs, or autophagy machinery to remove the damaged membranes. Atg8ylation is potentially a cellular defence mechanism against membrane damage. Our data suggest that Atg8ylation protects cells from cell death by selectively removing the damaged lysosomes. Atg8ylation generally involves recruitment of LC3B and lipidation of LC3B and GABARAP, however, LC3 subfamily of proteins is largely dispensable for Atg8ylation and related processes(Goodwin *et al*., 2021; Kaur *et al*, 2023; Lee *et al*, 2020; Nakamura *et al*., 2020). In contrast, ORF3a induced Atg8ylation involves both LC3 and GABARAP subfamily of proteins.

Previous report suggest that SARS-CoV-2 protein ORF3a induces lysosomal damage and inhibits fusion of autophagosomes with lysosomes by interfering with the autophagy related SNARE complexes (Miao *et al*., 2021; Walia *et al*., 2024). While our studies confirm the role of ORF3a in inducing lysosomal damage, we demonstrate the mATG8s get recruited to damaged lysosomes and facilitate the selective lysophagy of damaged lysosomes by recruiting lysophagy receptor TRIM16 and its partner Gal3 to damaged lysosomes. Our data showing role of ORF3a on mTOR inhibition also suggest lysophagy induction. Overall, our findings extend the role of ORF3a in induction of lysophagy in addition to its previously established roles in autophagy and lysosomal pathway.

Recent studies have shown TECPR1 as a new regulator of Atg8ylation. TECPR1controls LC3B and GABARAP lipidation during lysosomal stress(Boyle *et al*, 2023; Corkery *et al*, 2023; Kaur *et al*., 2023). Interestingly, TECPR1 together with ATG16L1 conjugates ATG8s to membranes. TECPR1 binds to ATG5-ATG12 via an ATG5 interacting region and it possesses an LIR for ATG8 binding(Wang Y, 2023). While TECPR1 affects ATG8 conjugation and potentially recruits ATG8s to damaged lysosomes, it remains elusive whether TECPR1 or other potential E3 like proteins can form complexes with ATG5-12 system and regulate Atg8ylation during starvation induced autophagy. Our data showing the role of ATG16L1 alone suggest that ORF3a induced Atg8ylation may be independent of TECPR1, however, future studies are needed to investigate if lysosomal damage induced by SARS-CoV-2 infection or ORF3a require combined action of ATG16L1 and TECPR1.

Overall, our findings elucidate the mechanism by which the SARS-CoV-2 protein ORF3a induces lysosomal membrane Atg8ylation and establish mATG8s and the ATG8 conjugation machinery as critical regulators of this response. These findings expand our understanding of how SARS-CoV-2 perturbs the endolysosomal system and reveal Atg8ylation as an important cellular response to viral induced lysosomal stress. Given the emerging role of Atg8ylation in membrane damage, repair, and pathogen host interactions, this pathway may represent a potential therapeutic target in infectious diseases associated with endolysosomal dysfunction or exploitation of the autophagy machinery.

## METHODS

### Antibodies and reagents

The following antibodies and dilutions were used: Flag (mouse monoclonal Sigma; F1804, used at 0.5 µg/ml and 1:1,000 for (WB); 1:250 (IF)); GFP (mouse Santa Cruze Biotechnology: sc-9996; 0.5 µg/ml IP and 1:1,000 (WB)); Galectin3 (mouse Santa Cruze Biotechnology: sc-32790; 1:200 (IF)); LAMP2 (mouse Santa Cruze Biotechnology: sc-19991; 1:200 (IF)); LC3 (rabbit; MBL International PM036, 1:500 (IF)); FIP200 (rabbit; Proteintech; 17250-1-AP, 1:200 (IF)); FIP200 (rabbit Cell Signaling Technology; 12436, 1:1000 (WB)); mTOR (rabbit Cell Signaling Technology; 2983, 1:200 (IF)); TFEB (rabbit Cell Signaling Technology; 4240, 1:200 (IF)); LAMP1 (rabbit Cell Signaling Technology; 9091, 1:200 (IF)); pP70S6K (rabbit Cell Signaling Technology; 9234, 1:1000 (WB)); P70S6K (rabbit Cell Signaling Technology; 9202, 1:1000 (WB)); mCherry (rabbit Cell Signaling Technology; 43590, 1:1000 (WB)); HA (rabbit Cell Signaling Technology; 3724, 1:100 (IP) 1:1000 (WB)); LC3B (Cell Signaling Technology; 3868, 1:400 (IF)), LC3C (rabbit Cell Signaling Technology; 14736, 1:400(IF)), GABARAP ((rabbit Cell Signaling Technology; 13733, 1:400(IF)), ALIX (rabbit Cell Signaling Technology; 92880, 1:200 (IF)); ATG16L1 (rabbit; MBL International PM040, 1:1000 (WB)); Dynabeads Protein G (Thermo Fisher Scientific 10003D 50µl/ IP); Bafilomycin A1 (Baf A1, InvivoGen, 13D02-MM); Lipofectamine 2000, Thermo Scientific, 11668019; anti-rabbit Alexa Fluor-488 (A-11034), -568 (A-11036) and -647 (A-21245), and anti-mouse Alexa Fluor -488 (A-11029), -568 (A-11004) and -647 (A-21235) (1:500 for IF) were from ThermoFisher Scientific.

### Cell culture

HEK 293T and HeLa cells were maintained in ATCC recommended media as described previously (Kumar *et al*., 2020a). Hexa^KO^ and parental HeLa cells have been described previously and were maintained in DMEM supplemented with 10% fetal bovine serum and antibiotic as described previously (Nguyen *et al*., 2016). 3T3 cells were cultured in DMEM supplemented with 10% fetal bovine serum and antibiotic.

### Plasmids transfections

pDest-GFP-LC3A, pDest-GFP-LC3B, pDest-GFP-LC3C, pDest-GFP-GABARAP, pDest-GFP-GABARAPL1, pDest-GFP-GABARAPL2, GFP-TFEB and FLAG-TFEB have been described earlier(Kumar *et al*., 2020a). Plasmids related to SARS-CoV-2 ORF3a-mCherry (#165138), ORF3a-EGFP (#165121), NSP6-mCherry (#165133), NSP4-mCherry (#165132), NSP3-mCherry (#165131), GFP-Gal-3 (#73080), px459 sgFIP200 (#175024), px459 sgATG5 (#175023), PX458-sgTSC2 (Plasmid #128098) psPAX2 (#12260), pCMV-VSV-G (#8454), pRK5 Flag RagB (#112755), pRK5 GFP RagB Q99L(#112749), pLJM1 Flag Raptor (Plasmid #26633) and Flag pLJM1 RagB 99L (#19315) were acquired from Addgene. ORF3a-HA, ORF3aHA Y160A/V163G (Walia *et al*., 2024) and FLAG-TRIM16 have been described previously (Jena *et al*, 2018). Plasmids were transfected using Lipofectamine 2000 (Thermo Fisher Scientific) and polyethylenimine (PEI). Briefly, cells were seeded in different plates for experiments and plasmids were transfected with lipofectamine (1:1; 1μl lipofectamine for 1μg plasmid) or PEI (1:3; 1μg plasmid and 3μl PEI) after overnight transfection, cells were assayed for different experiments as explained in the figure legends.

### Generation of CRISPR mutant cells

ATG16L1 CRISPR in HeLa, HCT116 and 3T3 cells were generated as described earlier(Kumar *et al*, 2018). Briefly, the lentiviral vector carrying both Cas9 enzyme and a gRNA targeting ATG16L1 (GGTCACAAAGCTTAGTGCGC) were transfected into HEK293T cells together with the packaging plasmids psPAX2 and pCMV-VSV-G at the ratio of 5: 3: 2. Two days after transfection, the supernatant containing lentiviruses was collected and used to infect the cells. 36 hours after infection, the cells were treated with puromycin (1 mg/ml) for one week to select knockout cells. The knockouts were confirmed by western blotting. Hexa^KO^ LC3^TKO^, and GABARAP^TKO^ HeLa cells were described previously (Nguyen *et al*., 2016).

### Generation of ATG5 and FIP200 and TSC2 CRISPR mutant cells

Px459 sgFIP200, PX458 sgTSC2 and PX459 sgATG5 were obtained from Addgene. These gRNA in bicistronic Cas9/sgRNA mammalian expression vector are previously described(Assar & Tumbarello, 2020). The FIP200 gRNA 5′- TATGTATTTCTGGTTAACAC-3′ targeted exon 3, TSC2 gRNA 5′- TCTGCTGAAGGCCATCGTGC-3′ and the Atg5 gRNA 5′- AAGATGTGCTTCGAGATGTG-3′ targeted exon 2 were transfected into HeLa cells using PEI. To generate FIP200 and ATG5 knock-out cells, cells were seeded in 6- well plate and transfected with 2µg of each pX459 sgFIP200 and pX459 sgATG5 construct using PEI (ratio of 1:3). After 24h cells were washed with PBS and replenished with fresh complete media. Cells were selected using puromycin (1mg/ml) for one week. Knock-outs were confirmed by western blotting.

### High content microscopy

High content microscopy was carried out in 96 well plates as described (Kumar *et al*., 2021a). Briefly, cells in the 96 well plates were transfected wherever indicated in the figures, with plasmids. Following transfection, cells were fixed with 4% paraformaldehyde for 5 mins. Cells were permeabilized with 0.1% TritonX100 and blocked in 3% BSA for 30 mins followed by incubation with primary antibody for overnight and secondary antibody for 1 h. HCM analysis was carried out using a Cellomics HCS scanner and iDEV software (Thermo) as described (Kumar *et al*., 2021a).

### High content microscopy for Keima probes

HeLa WT Hexa^KO^ and ATG5^KO^ cells were plated in 96 well plates and transfected with TMEM192-Keima plasmids. Transfected cells were incubated with Hoechst 33342 for ten minutes in full media and then acquired for Keima fluorescence at 440nm and 560 nm using the Cell Insight CX7 High-Content Screening (HCS) Platform (Thermo)(Kumar *et al*, 2019).

### Immunofluorescence confocal microscopy

Immunofluorescence confocal microscopy was done using Stellaris confocal micrscope from Leica and CellInsight CX7 LZR pro. Briefly, cells were plated onto coverslips in 6 well or 12-well plates. Cells were transfected with plasmids wherever required. After transfected cells were fixed in 4% paraformaldehyde for 5 min. Cells were then permeabilized with 0.1% Triton X100 in 3% BSA, followed by blocking in 3% BSA. After blocking cells were incubated with primary antibodies for overnight, followed by washing with PBS three times. Cells were then incubated with appropriate AlexaFluor labelled secondary antibodies (Invitrogen) for 1 h at room temperature. Coverslips were then mounted using ProLong Gold Antifade Mountant (Invitrogen) and analyzed by confocal microscopy using confocal platform. Images were processed using Adobe Photoshop 22.5.1.

### Western blotting and co-immunoprecipitation assays

Western blotting and co-immunoprecipitation (co-IP) were performed as described previously(Kumar *et al*., 2018). For western blotting, cells were transfected with ORF3a or other plasmids as indicate in the figures. Following transfection, cells were lysed using RIPA buffer (Tris-Hcl 25mM (pH 7.4), NaCl 150mM, sodium deoxycholate 0.5%, SDS 1%, NP40 1%) containing protease inhibitor cocktail, for 45 min at 4°C, followed by centrifugation at 12000xg for 15 min. Supernatant was boiled in samples buffer for 5 min and subjected to SDS-PAGE for separation, followed by transferring the proteins to PVDF membranes. Membranes were incubated with primary antibodies for overnight, followed by three washings and subsequent incubation with HRP-labelled secondary antibodies. Blots were detected by enhanced chemiluminescence using ChemiDoc MP.

For co-IP, cells were lysed in NP-40 buffer containing protease inhibitor cocktail (Roche, cat# 11697498001) and PMSF (Sigma, cat# 93482). 10% of the lysates were saved for inputs, rest of the lysates were incubated with 5 µg antibody at 4°C for overnight. This was followed by incubation of the antibody lysate mixture with Dynabeads protein G (Life Technologies) for 4h at 4°C. Lystaes with antibody-bead mixture were washed three times with PBS followed by boiling in SDS containing loading buffer. Supernatant was collected and run on SDS-PAGE for immunoblotting which was done as explained above. Densitometric analysis was carried out using the ImageJ software.

### PI incorporation assay using HCM

HeLa^WT^ or LC3^TKO^, GABARAP^TKO^ and ATG16L1^KO^, ATG5^KO^, and Hexa^KO^ cells were transfected with plasmids using PEI as indicated in the figures. Transfected cells were washed with PBS and incubated with PI (1:1000) for 15 min. Colocalization between PI and Hoechst was quantified using HCM.

### Cell proliferation assay

For cell proliferation assay, cells were seeded in 96 well plates and transfected with plasmids as indicated in figures. After overnight transfection, cells were incubated with (3-(4,5, -dimethylthiazole-2-yl) -2,5-diphenyltetrazolium bromide (MTT)) dye for 4h, the formazan crystals were dissolved in 100ul DMSO and optical density (OD) was measured at 570 using multimode plate reader(Kumar *et al*, 2013).

### RNA-seq

For RNAseq, cells were transfected with ORF3a-mCherry. After overnight transfection, RNA was extracted using Trizol reagent (Invitrogen, CA, USA) and RNA was quantified using Nanodrop 2000 (Thermofisher Scientific, Massachusetts, USA). Before, processing the samples for RNAseq, the integrity of RNA was evaluated on 1% agarose gel (Lonza, Belgium). For Illumina sequencing of RNA profile, KAPA RNA HyperPrep Kit was used. Briefly, Poly(A) RNA was purified from total RNA (5µg) using poly-T oligo-attached magnetic beads using two rounds of purification. Following purification, the mRNA was fragmented into small pieces using divalent cations (magnesium) under elevated temperature. Then the cleaved RNA fragments were reverse transcribed to create the final cDNA library using random priming. The average insert size for the paired-end libraries was 300 bp (±50 bp). dscDNA was prepared by combined second strand synthesis and A-tailing. Following adapter ligation and library preparation, the paired-end sequencing was carried out on an Illumina NovaseqTM 6000 at the (LC Sciences,USA) following the manufacturer’s recommended protocol. Using the Illumina paired-end RNA-seq approach, the transcriptome was sequenced, generating a total of 2 × 150 million bp paired-end reads. Raw reads were checked for base quality and adapter content using FastQC (v0.11.9). Fastp(v0.12.4) was used to remove adapter content and trim low-quality bases. Hisat2 (v0.12.0) was used to align reads to the genome of Homo sapiens(hg38). Feature Counts (v1.22.0) was used to generate gene level counts for differential gene expression using DESeq2 (v1.34.0). Genes with FDR < 0.05 after applying wald’s test were considered differentially expressed (DE) genes. DE genes were used for Gene Ontology and KEGG enrichment using the R package clusterProfiler (v4.12.0) and plotted using enrichplot. Genes with log2 fold change value below 0 are downregulated genes whereas genes with log2 fold change value above 0 are upregulated genes. An FDR cutoff of 0.05 is used to identify differentially expressed genes.

### Statistical analyses

Data are expressed as means ± SEM (n ≥ 3). Data were analyzed with a paired two-tailed Student’s *t-*test or analysis of variance (ANOVA) was used. Statistical significance was defined as † p ≥ 0.05, \**P* < 0.05, **p<0.01.

## Supporting information

Table S1 in the text

## Acknowledgments

We thank V. Deretic for sharing HeLa and 293T cell lines, and STX17 and GFP- mATG8 constructs, M. Lazarou for sharing HeLa^WT^, Hexa^KO^, GABARAP^TKO^ and LC3^TKO^ cells, S. Chauhan for FLAG-TRIM16 construct. This work was supported by ICMR grants IIRPIG-2024-01-00785, DBT grant BT/PR45023/COT/142/27/2022 and SERB grant SRG/2022/000085 to S.K.

---

**Figure S1.**
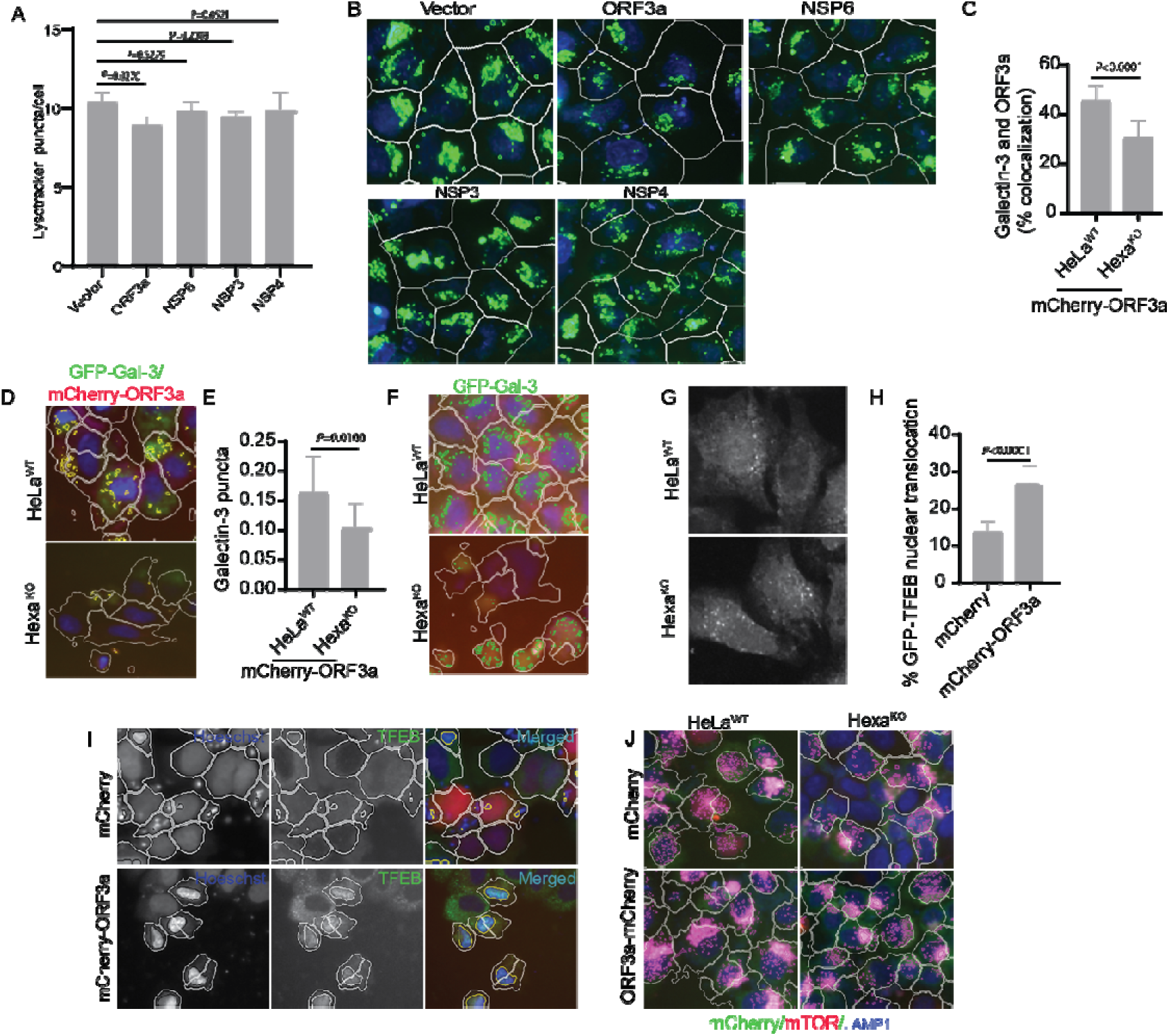
SARS-CoV-2 factor ORF3a but not NSP6s induce Atg8ylation. **(A, B)** HCM quantification and images showing the effect of expression of SARS-CoV-2 proteins ORF3a, NSP3, NSP4, NSP6 on lysotracker puncta formation. Masks; white: ORF3a, NSP3, NSP4 and NSP expressing cells, algorithm-defined cell boundaries; green masks: computer-identified lysotracker dots). p values were determine using GraphPad prism (n=9) ANOVA. Images, a detail from a large database of machine- collected and computer-processed images. Scale bar 10 µm. **(C, D)** HCM quantification and image analysis of the effect of Hexa^KO^ on ORF3a-mCherry and Gal3 colocalization. (>500 primary objects examined per well; minimum number of wells, 8). Masks; white: ORF3a-mCherry expressing cells, algorithm-defined cell boundaries; yellow masks: colocalization between Gal3 and ORF3a-mCherry dots). p values were determine using GraphPad prism (n=8) t-test. Scale bar 10 µm. **(E,F)** HCM quantification and image analysis of the effect of Hexa^KO^ on ORF3a induced GFP-Gal3 puncta formation. (>500 primary objects examined per well; minimum number of wells, 8). Masks; white: ORF3a-mCherry expressing cells, algorithm-defined cell boundaries; green masks: computer-identified GFP-Gal3 dots). p values were determine using GraphPad prism (n=8) t-test. Scale bar 10 µ **(G-I)** HCM quantifications and image analysis of the effect of ORF3a induced nuclear translocation of GFP-TFEB. (>500 primary objects/cells examined per well; minimum number of wells, 6). Masks; white: algorithm-defined cell boundaries; yellow outline: computer-identified colocalization between TFEB and Hoechst-33342 nuclear stain). p values were determine using GraphPad prism (n=6) t-test. Scale bar 10 µm **(J)** HCM images to test the effect of ORF3a-mCherry expression on colocalization between mTOR and LAMP1 in HeLa^WT^ and Hexa^KO^ cells. HCM, >500 cells counted per well; minimum number of valid wells 6, Masks; white: algorithm-defined cell boundaries; magenta: computer-identified co-localization between mTOR and LAMP2. Scale bar 5µm.

**Figure S2.**
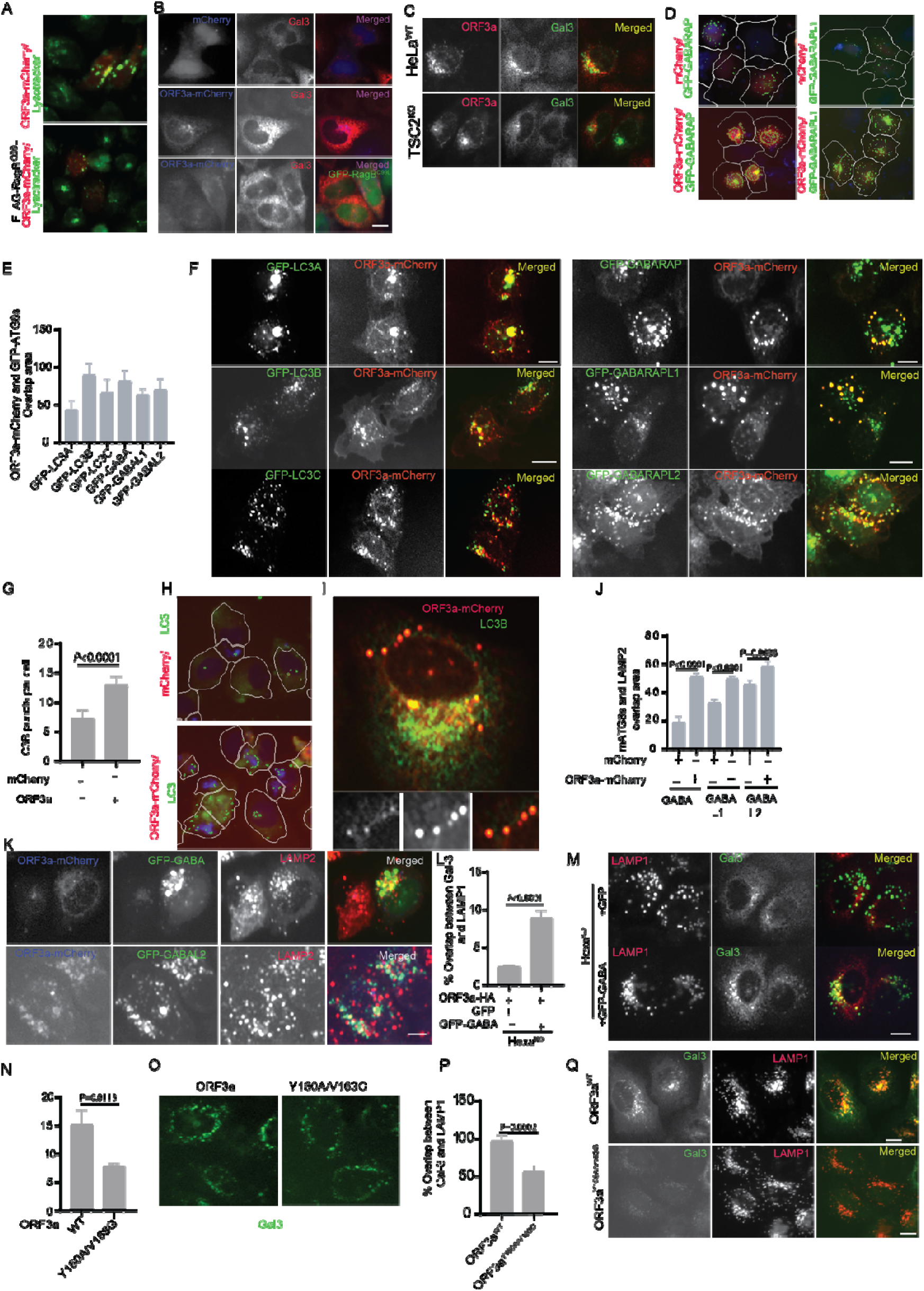
ORF3a results in mATG8 puncta formation. **(A)** Micrograph showing the effect of RagBQ99L expression on orf3a mediated lysotracker puncta formation. Scale bar 5µm. **(B)** Micrographs showing the effect of RagBQ99L expression on ORF3a mediated Gal3 puncta formation. Scale bar 5 µm. **(C)** Micrograph showing the effect of ORF3a expression on Gal3 puncta formation in Hela^WT^ and TSC2^KO^ cells. Scale bar 5µm. **(D)** HCM imaging showing the effect of ORF3a-mCherry or control vector on formation of GFP-GABARAP and GFP-GABARAPL1 puncta. **(E,F)** HCM quantification and micrograph analysis of colocalization between ORF3a- mCherry and GFP-ATG8. **(G,H)** HCM quantifications and imaging showing the effect of ORF3a expression on endogenous LC3 puncta formation. Masks; white: ORF3a, expressing cells, algorithm-defined cell boundaries; green masks: computer-identified LC3 dots). p values were determine using GraphPad prism (n=9) t-test. Scale bar 10 µm. **(I)** Confocal image analysis of colocalization between ORF3a- mCherry and endogenous LC3B . **(J,K)** HCM quantifications and image analysis showing the effect of ORF3a expression on localization of GABARAPs (GFP-GABARAP, GABARAP L1, GABARAP L2) to lysosomes. p values were determined using GraphPad prism (n=5) t-test. Pseudo colours: Red, LAMP2; green, GFP- mATG8s; blue, ORF3a-mCherry. p values, (n=5) ANOVA. Scale bar 5 µm. **(L,M)** HCM quantification and image analysis of complementation of Hexa^KO^ cells with GFP-GABARAP on colocalization between Gal-3 and LAMP1 upon ORF3a expresion. **N,O)** HCM quantifications and image analysis showing the effect of ORF3a-HA or ORF3aY160A/V163G-HA expression on Gal3 puncta formation. p values were determine using GraphPad prism (n=9) t-test. Scale bar 5 µm. **(P,Q)** HCM quantifications and image analysis showing the effect of ORF3a-HA or ORF3aY160A/V163G-HA expression on Gal3 and LAMP1 colocalization in HeLa cells. p values were determine using GraphPad prism (n=9) t-test. Scale bar 5 µm.

**Figure S3.**
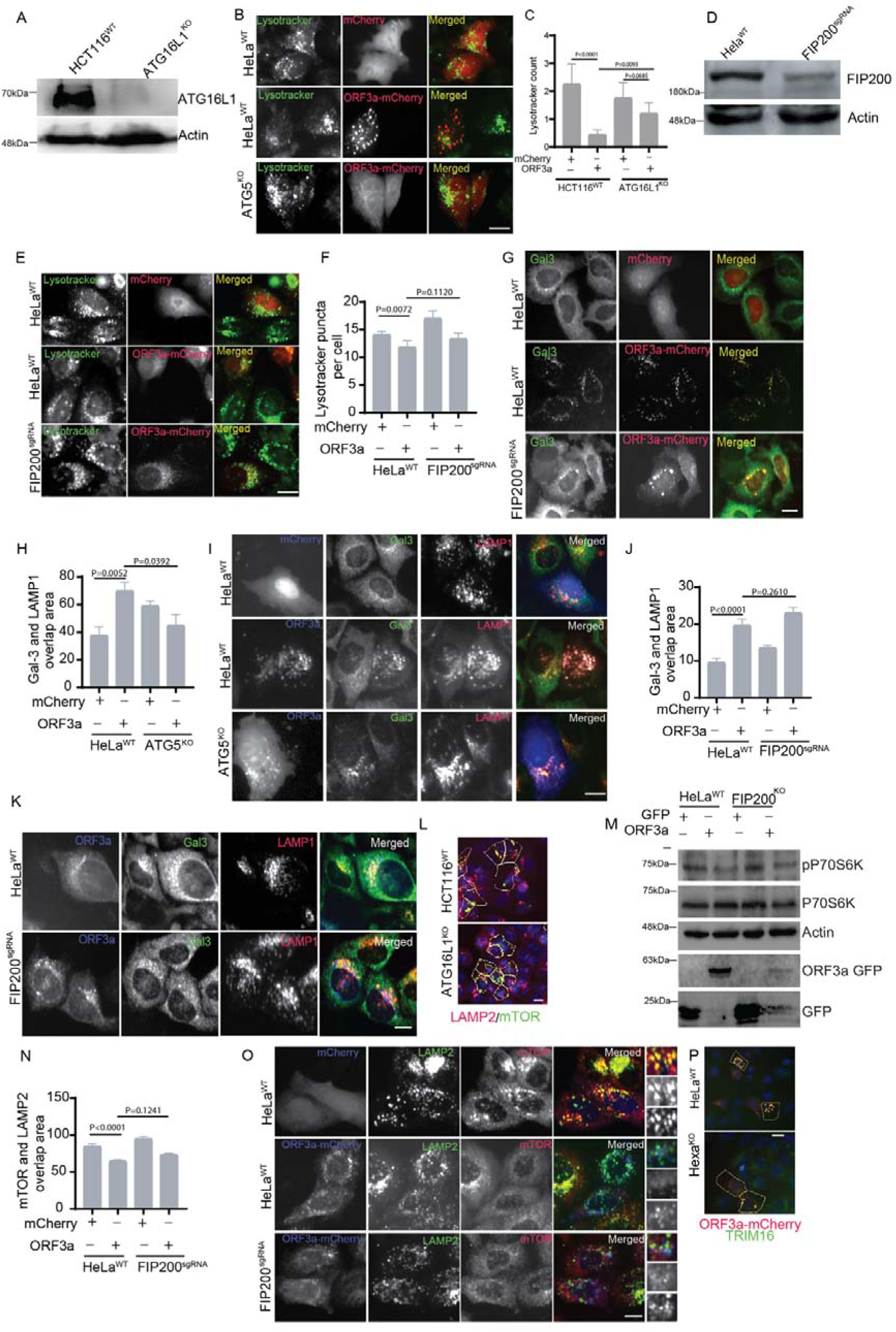
Atg8ylation is independent of FIP200. **(A)** Western blot showing ATG16L1 knock out (ATG16L1^KO^) in HCT116 cells. **(B,C)** HCM quantification and image analysis of effect of ORF3a expression on lysotracker puncta formation in HCT116^WT^ and ATG16L1^KO^ cells. P values were determined using GraphPad prim (n=6) ANOVA. Scale bar 5µm. **(D)** Western blot showing FIP200 downregulation in FIP200 CRISPR cells. **(E,F)** HCM quantification and micrograph analysis of the effect of ORF3a expression on lysotracker puncta formation in Hela^WT^ and FIP200^KO^ cells. Scale bar 5µm **(G)** Micrograph showing the effect of ORF3a expression on Gal3 puncta formation in HeLa^WT^ and FIP200^KO^ cells. Scale bar 5µm. **(H,I)** HCM quantification analysis of the effect of ATG5^KO^ on colocalization between Gal3 and LAMP1 in HeLa cells expressing mCherry or ORF3a-mCherry. Micrograph showing the effect of ATG5^KO^ on colocalization between Gal3 and LAMP2 in HeLa cells expressing mCherry or ORF3a-mCherry. Scale bar 5 µm. **(J,K)** HCM quantifications to analyze the effect of FIP200^KO^ on colocalization between Gal3 and LAMP2 in HeLa cells expressing mCherry or ORF3a-mCherry. Micrograph showing the effect of FIP200^KO^ on colocalization between Gal3 and LAMP2 in HeLa cells expressing mCherry or ORF3a-mCherry. Scale bar 5 µm. **(L**) HCM image analysis of the effect of ATG16L1^KO^ on ORF3a induced colocalization between LAMP2 and mTOR in HCT116 cells. p values were determined using GraphPad prism (n=5) ANOVA. Masks; white: ORF3a expressing cells, algorithm defined boundaries. Scale bar 5µm. **(M)** Immunoblot analysis of the effect of FIP200^KO^ on ORF3a induced phosphorylation of P70S6K. **(N,O)** HCM quantifications to analyze the effect of FIP200^KO^ on colocalization between mTOR and LAMP2 in HeLa cells expressing mCherry or ORF3a-mCherry. p values were determined using GraphPad prism (n=9) t-test. Scale bar 5 µm. **(P)** HCM imaging showing colocalization of ORF3a-mCherry with FLAG-TRIM16 in Hela wild type and Hexa^KO^ cells. Scale bar 5 µm .

**Figure S4.**
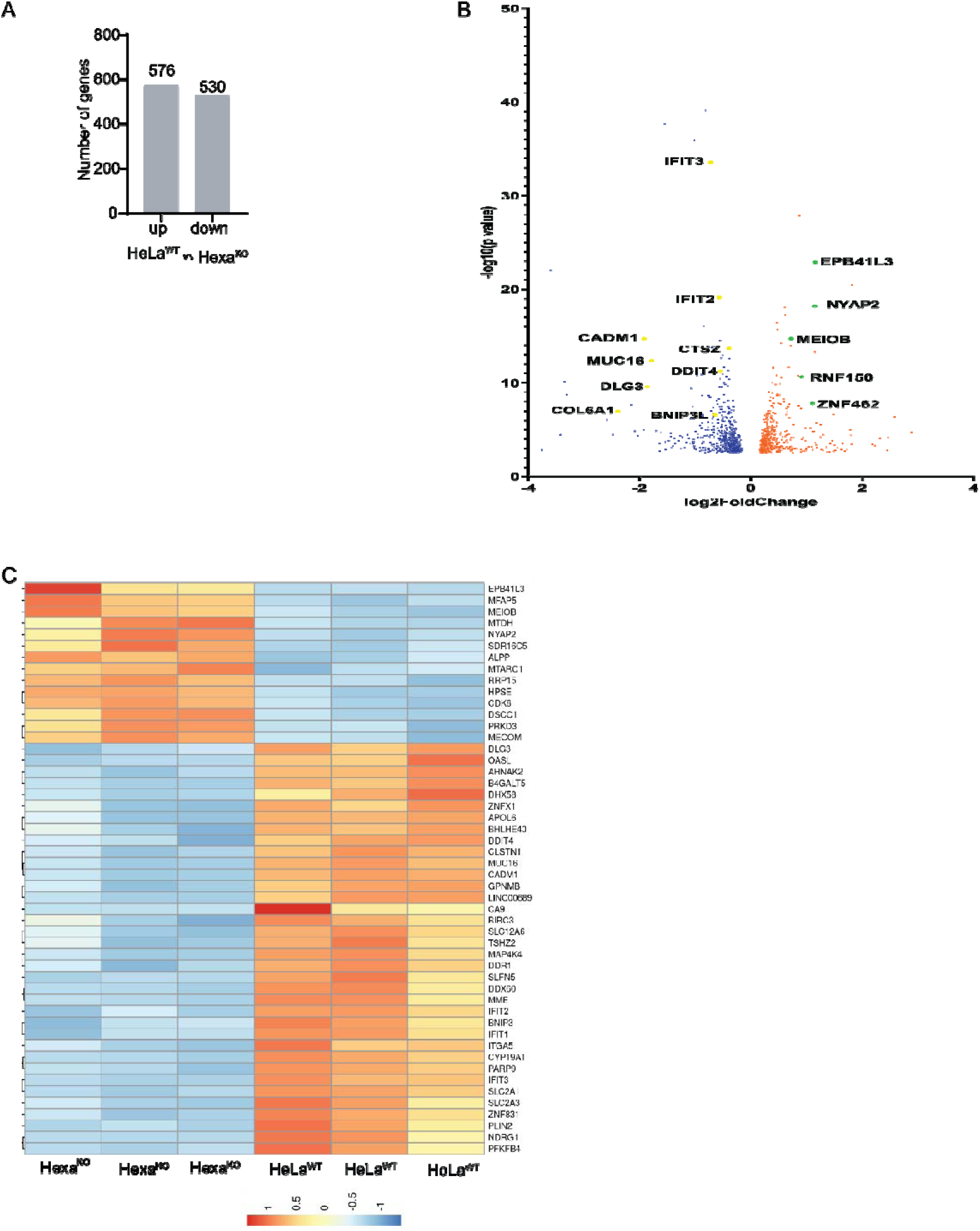
STX17 and mATG8s control expression of genes involved in important biological function. **(A)** Total number of genes those were upregulated or downregulated in Hexa^KO^ or. HeLa^WT^, Hexa^KO^ cells were transfected with ORF3a, RNA was isolated and RNAseq was performed as described in materials and methods. **(B)** Volcano plot showing the effect of Hexa^KO^ on differential gene expression. Yellow circles indicate genes downregulated in Hexa^KO^ cells; green circles genes upregulated in HexaKO cells. Cut-off used in the plot (P < 0.05). **(C)** Heat maps showing top 50 genes those were upregulated or downregulated in Hexa^KO^ (E) .

**Figure S5.**
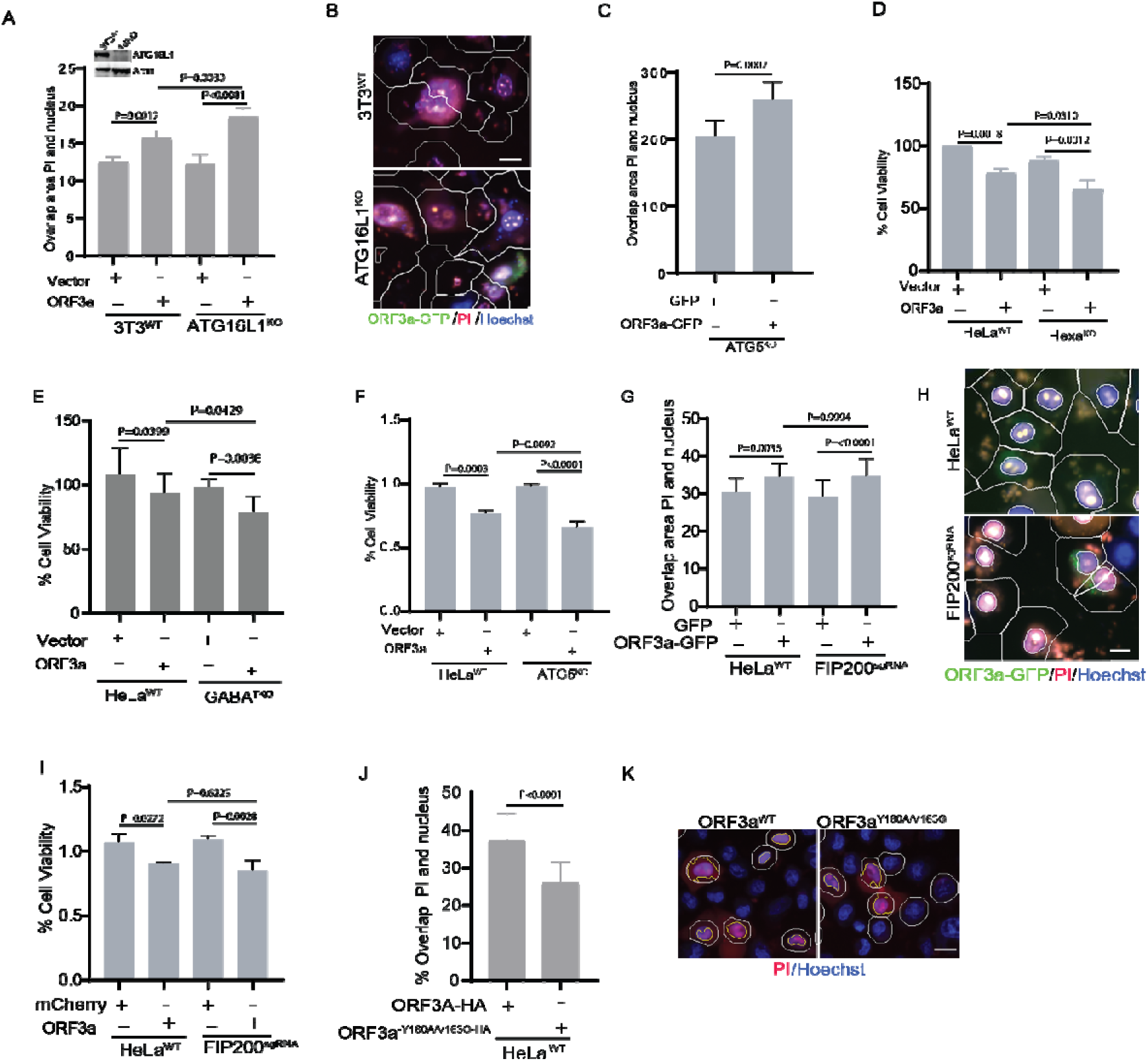
FIP200 is not involved in ORF3a induced cell death. (A,B) WT and ATG16L1^KO^ 3T3 cells were transfected with GFP or ORF3a-GFP, cells were stained with PI and analyzed under HCM for PI incorporation assay. Colocalization of PI and Hoechst stain was quantified in GFP or ORF3a-GFP transfected cells. Masks; white: GFP or ORF3a-GFP expressing cells, algorithm-defined cell boundaries. p values were determine using GraphPad prism (n=9) ANOVA. Scale bar 10 µm **(C)** HCM quantification analysis of the effect of ORF3a induced cell death in ATG5^KO^ cells which was marked by PI uptake. **(D)** WT and Hexa^KO^ HeLa cells were transfected with mCherry or ORF3a-mCherry, after overnight transfection, cells were incubated with MTT dye for 4h and absorbance was measured at 450nm. p values were determine using GraphPad prism (n=16) ANOVA. **(E)** WT and GABA^TKO^ HeLa cells were transfected with mCherry or ORF3a-mCherry, after overnight transfection, cells were incubated with MTT dye for 4h and absorbance was measured at 450nm. P values were determine using GraphPad prism (n=16) ANOVA. **(F)** WT and ATG5^KO^ HeLa cells were transfected with mCherry or ORF3a-mCherry, after overnight transfection, cells were incubated with MTT dye for 4h and absorbance was measured at 450nm. p values were determine using GraphPad prism (n=16) ANOVA. **(G,H)** WT and FIP200^KO^ HeLa cells were transfected with GFP or ORF3a- GFP, cells were stained with PI and analyzed under HCM for PI incorporation assay. Colocalization of PI and Hoechst stain was quantified in GFP or ORF3a-GFP transfected cells. Masks; white: GFP or ORF3a-GFP expressing cells, algorithm-defined cell boundaries. p values were determine using GraphPad prism (n=9) ANOVA. Scale bar 10 µm. **(I)** WT and FIP200^KO^ HeLa cells were transfected with mCherry or ORF3a-mCherry, after overnight transfection, cells were incubated with MTT dye for 4h and absorbance was measured at 450nm. p values were determine using GraphPad prism (n=16) ANOVA. **(J,K)** HCM quantification and HCM analysis of effect of ORF3a-HA and ORF3a-Y160A/V163G-HA expression on PI uptake in HeLa cells.

**Table S1: RNAseq analysis of the effect of Hexa^KO^ on global gene expression in ORF3a expressing cells.** HexaKO and parental HeLa cells were transfected with ORF3a-GFP. Experiment was terminated when >85 % cells were ORF3a-GFP positive. Total RNA was isolated from the cells and RNAseq was carried out using Illumina paired-end RNA-seq approach as explained in methods.

